# Boreal and subarctic freshwaters harbour a diversity of jumbophages

**DOI:** 10.64898/2026.04.22.720137

**Authors:** Meeri Niemi, Karyna Karneyeva, Lotta-Riina Sundberg, Hanna M. Oksanen, Elina Laanto

**Author notes:** corresponding author Address: Department of Biological and Environmental Science and Nanoscience Center, PO Box 35, FI-40014 University of Jyväskylä, Finland.

## Abstract

Bacteriophages (phages) are major drivers of microbial evolution and ecology, yet their diversity and functional roles remain poorly characterised in many natural environments, such as in freshwater systems. In boreal and subarctic freshwater habitats, where bacteria are typically slow-growing and nutrient-limited, phages are predicted to have a critical role in host regulation and horizontal gene exchange. However, only a few isolates have been obtained from such environments, leaving the genetic and functional diversity of these phages largely unexplored. Here, we present a collection of 40 bacteriophages isolated from boreal lakes and rivers using a set of diverse freshwater bacterial hosts. Despite using conventional isolation methods, eight of the isolates possess genomes larger than 200 kilobases and are classified as jumbophages. All jumbophages exhibited myovirus morphology and comparatively slow infection dynamics. These jumbophages include the first known representatives infecting members of *Janthinobacterium* and *Herbaspirillum*. Comparative genomic and phylogenetic analyses show that nearly all genomes are distinct from previously described phages, indicating substantial novelty. Diverse auxiliary metabolic and anti-defence systems were identified, including putative NAD^+^ salvage and acyl carrier protein modules, along with predicted Anti-Thoeris and Anti-CBASS elements. The *Pseudomonas-*infecting jumbophage Ahti encoded homologues of all 21 core genes that define the nucleus-forming family *Chimalliviridae*. Additionally, Ahti displayed compartmentalisation of DNA during infection, establishing it as the first freshwater nucleus-forming phage. These findings expand our understanding of the ecological, genomic, and functional diversity of phages in boreal environments, and highlight the role of freshwater ecosystems as significant reservoirs of novel viral lineages.

**Importance:** Bacteriophages are viruses that infect bacteria and play important roles in shaping microbial communities and nutrient cycling in natural waters. However, much of what is known about their diversity comes from sequencing data alone, without environmental isolates that allow direct studies on their biology. In this study, we described 40 bacteriophages from boreal and subarctic lakes and rivers using freshwater bacterial hosts, including species of *Flavobacterium, Pseudomonas, Janthinobacterium* and other lesser-known phage hosts. Eight of these viruses had exceptionally large genomes, categorising them as jumbophages. These isolates included the first known jumbophages infecting *Janthinobacterium* and *Herbaspirillum*, as well as the first freshwater jumbophage shown to form a nucleus-like structure during infection. Together, these findings reveal boreal freshwaters as an important source of previously unknown virus diversity and provide new model systems for exploring phage biology in environmentally relevant contexts.

## Introduction

The boreal biome, the world’s largest terrestrial biome, contains extensive networks of lakes, rivers, and wetlands that support diverse microbial communities adapted to cold, nutrient-poor, and humic-rich conditions (Berggren et al., 2010; Grosbois et al., 2023). These ecosystems are highly vulnerable to rapid environmental change, including warming-driven shifts in water temperature, reduced ice cover duration, altered mixing regimes, and increasing terrestrial organic matter inputs that drive freshwater browning. Together, these changes can reshape habitat structure, nutrient availability, and microbial community composition in northern freshwaters (Woolway et al., 2020; Blanchet et al., 2022). The functioning of these ecosystems is strongly influenced by bacteriophages (phages), which regulate their bacterial host populations, mediate horizontal gene transfer, and reprogram host metabolism (Naureen et al., 2020). In boreal freshwaters, where microbes are typically slow-growing and highly specialised (Crevecoeur et al., 2022), phages represent a major genetic reservoir. Large phages, in particular, are predicted to encode novel proteins and unique functions (Al-Shayeb et al., 2020). Despite this, environmental phages remain poorly characterised, with most current insights derived from metagenomic surveys (Hayes et al., 2017). While metagenomic approaches reveal extensive viral diversity, they are limited by challenges in assembly and host assignment. Isolate-based studies are thus essential as they provide direct evidence of phage– host interactions and enable the discovery of novel virus types (Laanto et al., 2017). Isolating unique phages not only expands our understanding of viral diversity but also allows sequence-based predictions to be linked to observable infection phenotypes, clarifying the molecular basis of phage–host interactions and their evolutionary trajectories within these complex freshwater systems.

Among such unique isolates, jumbophages are non-model bacteriophages distinguished by their unusually large genomes exceeding 200 kilobases (Hendrix, 2009), which encode diverse and poorly characterised functional repertoires, making them key targets for isolate-based exploration. Jumbophages have been isolated from different environments, including soil (Kozlova et al., 2024) and aquatic systems (Misol et al., 2020), suggesting a broad ecological distribution and prevalence. These phages exhibit several distinct biological features that set them apart from smaller, more typical and well-characterised phages. For example, some jumbophages form a proteinaceous “phage nucleus” during infection, a compartmentalised structure that physically separates phage DNA from host cytoplasm, protecting replication processes from bacterial defence systems (Chaikeeratisak et al., 2017; Laughlin et al., 2022). Many jumbophages also employ novel anti-defence strategies (van den Berg et al., 2023; Tesson et al., 2024). In addition, jumbophage genomes can harbour auxiliary metabolic genes (AMGs) that modulate host metabolism to favour phage replication (Muscatt et al., 2023), as well as extensive arrays of tRNA genes that may optimise translation of phage proteins (Yoshikawa et al., 2018). These features highlight jumbophages as complex viruses that offer valuable insight into genome size evolution and phage–host interactions.

Despite the growing recognition of jumbophages as ecologically and evolutionarily significant, studies based on isolated phage–host systems remain limited, possibly due to the technical challenges and time-intensive nature of their isolation and characterisation. Much of our current knowledge is based on metagenomic data (Iyer et al., 2021), leaving substantial gaps in our understanding of jumbophage biology, diversity, and host interactions. In this study, we address this gap by characterising a unique collection of phages isolated using multiple bacterial hosts from boreal and subarctic freshwater environments, both of which are largely unexplored in phage research. Remarkably, a significant portion of the collection (8 out of 40) were jumbophages. Five of these jumbophages infect *Flavobacterium* hosts, suggesting that members of this genus may be suitable for large-phage replication. Notably, the collection also includes the first reported jumbophages infecting members of *Herbaspirillum* and *Janthinobacterium*, two bacterial genera for which phage–host interactions remain largely unknown. Our study also reports the first nucleus-forming jumbophage isolated from freshwater. Together, these findings expand the known host range and ecological scope of jumbophages, providing valuable isolate-based data to advance the phage field.

## Materials and Methods

### Environmental sampling and culture conditions

Freshwater samples (50 mL–2 L) were collected from surface waters of lakes and rivers across Finland (Fig. 1). These samples were used for the isolation of bacteria and their associated phages (see Phage isolation). All bacterial and phage cultures were grown in one-fifth diluted Luria–Bertani medium (1/5 LB) prepared in tap water. Solid and top-agar layers initially contained 1 % and 0.5–0.7 % (w/v) agar, respectively. To improve plaque visibility, top agar was later replaced with 0.5 % (w/v) low-melting agarose. Melted top agar was maintained at 47 °C (agar) or 36 °C (low-melt agarose) prior to plating. Unless otherwise stated, cultures were incubated aerobically at room temperature (RT; 22 °C) with shaking (120 rpm).

**Figure 1.**
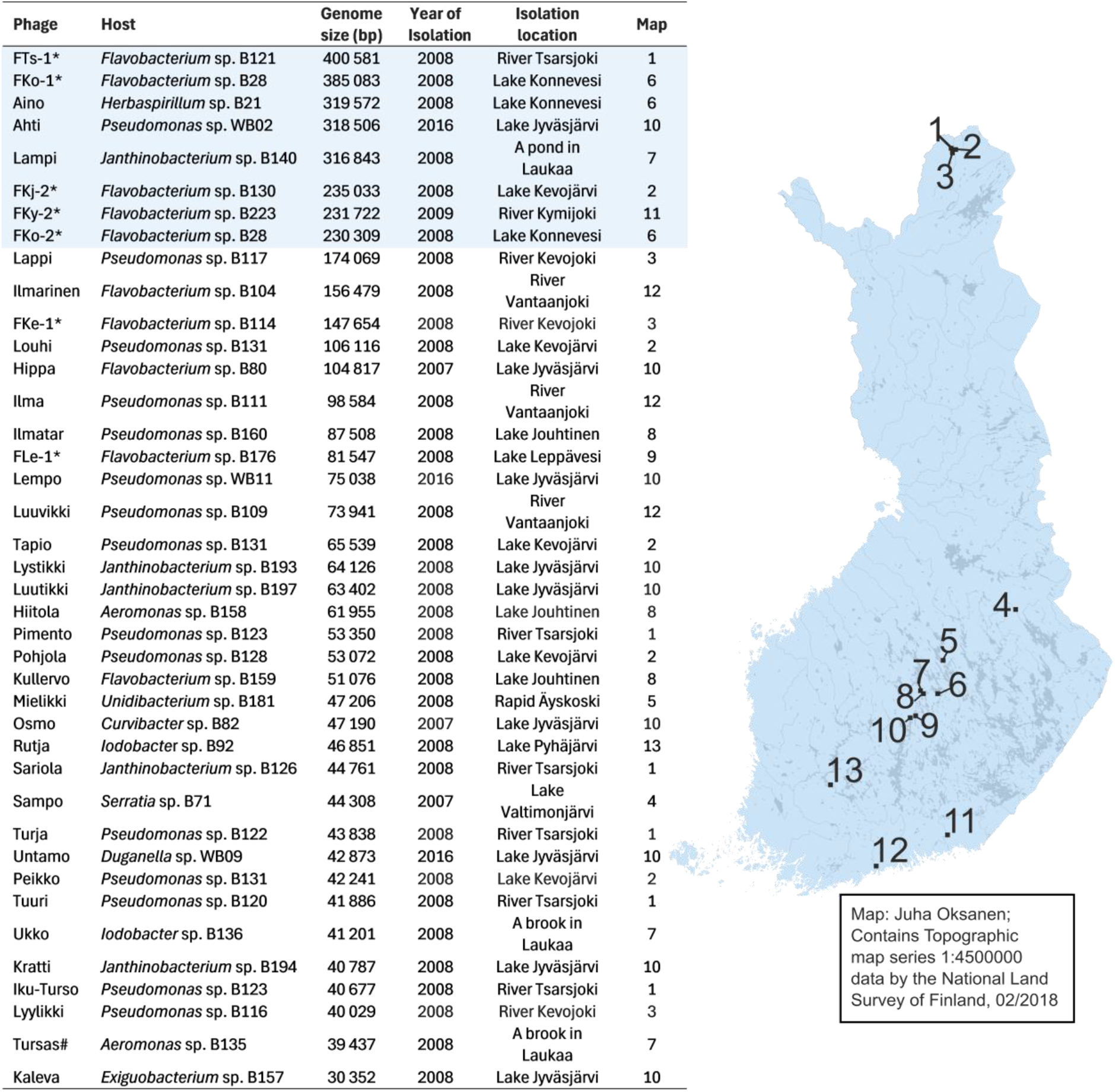
Collection of freshwater phages and their bacterial hosts. The phages were isolated from Finnish rivers and lakes (labelled 1–13, Table S1) in this study, except for those marked with an asterisk (Laanto et al., 2011) and a hash sign (Almeida et al., 2019). Year of isolation refers to the isolation of the phage. The jumbophage isolates are highlighted in blue.

### Bacterial host isolation and identification

Bacterial strains used as phage hosts are listed in Figure 1 and Supplementary Table S1. New bacterial isolates were obtained by plating 100 µL of freshwater samples onto 1/5 LB agar and incubating for two days (RT). Single colonies were purified by forming at least three consecutive pure cultures. Host taxonomy was assigned at genus level using a 95 % similarity threshold based on 16S rRNA sequence similarity to reference sequences from the NCBI database (Supplementary material, Fig. S1).

### Phage isolation and purification

Phage isolation was performed using enrichment cultures and bacterial hosts obtained from the same water source; in some cases, hosts originated from earlier sample collections at the same locations. For enrichment, water samples were filtered through 0.45 µm PES syringe filters (Sarstedt), and LB medium was added to achieve a final 1/5 LB concentration. Overnight grown bacterial cultures were diluted tenfold into the filtrate and incubated with shaking (RT, 120 rpm) until the culture was turbid (1–2 days). Aliquots (300 µL) of the enrichment cultures were plated using the plaque assay method: 300 µL of overnight grown host culture, phage sample (diluted as necessary), and 3 mL of molten top agar were poured onto 1/5 LB agar plates and incubated for 48 h. Individual plaques were picked into 500 µL of 1/5 LB and purified through three successive rounds of single-plaque assay.

High titre phage lysates (Table S2) were prepared from semi-confluent or confluent plates by collecting top agar into 4 mL of medium per plate, shaking for 4 h (RT), centrifuging (8000 × *g*, 10 min, 4 °C), and filtering (0.22 or 0.45 µm PES). Alternatively, confluent plates were overlaid with 5 mL of medium and shaken for 4–6 h (6 °C, 90 rpm) before liquid collection, centrifugation (4000 × *g*, 15 min), and filtration. Jumbophage particles were further purified by PEG precipitation followed by sucrose-gradient ultracentrifugation, negatively stained, and imaged using transmission electron microscopy (Supplementary material). Lysates were stored at 4 °C for short-term use or at –80 °C with 20 % (v/v) glycerol for long-term storage.

### Phage genome extraction, sequencing and assembly

Phage genomic DNA was extracted from filtered lysates according to the method of Santos (1991) with modifications described by Laanto and Oksanen (2023). Sequencing was performed by Novogene using the Illumina NovaSeq platform with paired-end 150 bp reads. Raw reads were uploaded to the Phagenomics phage genome analysis portal (www.phagenomics.net; accessed in February 2025) for assembly and annotation. Reads were also assembled with Velvet (version 1.2.10) in the Geneious platform (Biomatters Ltd) for comparison. When automated genome assembly by the Phagenomics portal failed, manual assemblies were conducted using the Galaxy platform (The Galaxy Community, 2024, Supplementary material). Final assemblies were re-uploaded to Phagenomics for downstream analyses.

### Genome annotation and comparative genomics

Genome annotation was performed via the Phagenomics portal against the PHROGs, pVOGs, COGs, and PHANNs databases (Table S3). Annotation summaries were visualised in R (v4.5.1, R Core Team, 2025). Jumbophages were additionally annotated using Pharokka and the Phold (Bouras et al., 2025) (Table S4). The closest reference genomes were identified against the INPHARED database (April 2025 release; Cook et al., 2021) by whole-genome BLAST (Camacho et al., 2009) for subsequent analysis with ViPTree (accessed in October 2025, Nishimura et al., 2017), VIRIDIC (accessed in September 2025, Moraru et al., 2020) and VirClust servers (accessed in October 2025, Moraru, 2023).

### Functional annotation and defence systems

Auxiliary metabolic genes (AMGs) in the jumbophage genomes were predicted using VIBRANT (v1.2.0) (Kieft et al., 2020). Annotated AMGs were manually checked against the annotated genomes and, in the case of rare AMGs, a more detailed analysis of the surrounding genes was performed. AMG island alignments were done using DiGAlign v2.0 (Nishimura et al., 2024) with Blastx. For the whole phage collection, anti-defence systems were identified with the AntiDefenseFinder mode of DefenseFinder (v2.0.2; Tesson et al., 2022), with additional searches performed for the jumbophages using DefenseFinder and PadLoc (v2.0.0; Payne et al., 2022). CRISPR arrays and *cas* genes were checked with CRISPRCasFinder (v1.1.2; Couvin et al., 2018). Putative homologues of the core jumbophage genes defined by Iyer et al. (2021) were identified in our genomes by searching for annotated matches. tRNA genes were extracted from all genomes using Biopython (Cock et al., 2009) in Python 3. Correlations between genome length and tRNA count were evaluated using Pearson’s and Spearman’s tests in RStudio (v2025.05.1). Plots were generated using ggplot2 (v3.5.2, Wickham, 2016) and dplyr (v1.1.4, Wickham et al., 2025).

### Jumbophage–host interactions

To measure the phage effect on host growth, all jumbophage hosts were grown overnight and diluted approximately 1:10 with medium to reach an OD_600_ of ∼0.1. Cultures were pipetted to a Honeycomb plate (100-well plate, Oy Growth Curves Ab Ltd) with 200 µL in each well. Phage was added at three multiplicities of infection (MOIs): 0.01, 0.1, and 1 to approximately 10^7^ CFU mL^−1^ of cells in the diluted culture using a volume of 10 µL. OD measurements were taken with Bioscreen C (Growth Curves Ltd) at 600 nm at 15-minute intervals for 25 hours without shaking. All treatments were performed in five replicates, and as controls, the OD of uninfected bacteria, medium, and phage lysate were also followed. To record plaque morphology, jumbophages were spotted on their respective isolation hosts using a series of dilutions to produce distinct plaques. Following 24–48 h of incubation, plates were imaged (ChemiDoc MP, Bio-Rad).

The host range of each jumbophage was tested against all jumbophage host strains (Table 1) using a double-agar overlay spot assay. Overnight cultures (300 µL) were mixed with 3 mL soft agar and poured onto agar plates. For initial screening, 10 µL of undiluted and 1:100 diluted phage lysates were spotted onto freshly prepared bacterial lawns. Sterile medium was used as a negative control, while the original host strains were used as the positive control. Plates were incubated (RT) and examined after 24 and 48 h for clear zones of lysis or plaque formation, which were recorded as evidence of infection. Bacteria–phage pairs showing infection were then retested in triplicate, using ten-fold serial dilutions spotted onto lawns and incubated for 24 h (RT).

### Phylogenetic analyses

A curated dataset of 105 large terminase subunit sequences (Table S5), combining our eight newly sequenced jumbophages with reference phages, was aligned with MAFFT (v7.526, Katoh & Standley, 2013) and analysed with IQ-TREE 2 (1,000 bootstraps, Minh et al., 2020) to infer phylogenetic relationships, with taxonomy assigned using ICTV/NCBI database and trees visualised in iTOL (v7.2.2, Letunic & Bork, 2024) (Supplementary material).

To assess the presence of nucleus-forming phage core genes in phage Ahti, a comparative sequence analysis was performed using the 21 core genes specific to chimallin-encoding phages as defined by Prichard et al. (2023). Protein sequences from nucleus-forming phages ΦKZ (NC_004629.1), vB_EcoM_Goslar (NC_048170.1), and vB_EamM_RAY (NC_041973.1) were used as reference sequences. The complete proteome of Ahti was queried against the core gene products using BLAST+ (blastp) with E-value ≤ 0.001, and similarity was assessed from bit scores.

### Detection of nucleus-like structures during infection

Bacterial cultures were infected, fixed, and DAPI stained (Supplementary material) to visualise DNA compartmentalisation. For imaging, 2 µL of sample was placed onto 1 % low-melting agarose pads and imaged by confocal microscopy. Phage nucleus formation was inferred from the appearance of centrally localised DNA focus in infected cells, in contrast to the diffuse DNA staining pattern observed in uninfected cells.

## Results

### Diverse freshwater phage–host systems recovered for genomic analysis

Across boreal and subarctic freshwater sites in Finland, the enrichment-based isolation strategy yielded 60 phage isolates, 40 of which were successfully sequenced (Fig. 1, Table S1). Sampling was conducted during the ice-free season, although a few phages were isolated from beneath the ice cover (e.g. FKo-1 and Hippa; Table S1). The genome-resolved phages were named using terminology from the Finnish national epic, the *Kalevala*, along with other names derived from Finnish mythology linked to aquatic environments. In total, 58 bacterial strains were isolated from freshwater samples (Table S1). Host identity was determined from partial 16S rRNA gene sequences for the 34 strains that served as hosts for phages (Fig. 1). Most isolates were Gram-negative freshwater bacteria, with the only Gram-positive strain belonging to *Exiguobacterium* sp., isolated from Lake Jyväsjärvi. Most isolates were assigned to *Pseudomonas* (15 strains) or *Flavobacterium* (10 strains), both common in freshwaters (Bernardet & Bowman, 2006; Pereira et al., 2018). Additional genera included *Janthinobacterium, Aeromonas, Undibacterium, Curvibacter*, and *Serratia*.

### Genome size variation highlights a subset of jumbophages

The largest number of collected phages was isolated with *Pseudomonas* (15 out of 40), including one jumbophage. Despite *Flavobacterium-*infecting phages accounting for a smaller portion of the collection (10 out of 40), half of all *Flavobacterium* phages had genomes exceeding 200 kb. The *Flavobacterium* jumbophages formed closely related branches on the proteomic tree (Fig. 2). To further investigate their relatedness, we identified shared protein clusters (PC) with VirClust. Protein cluster analysis (>30 % identity, >50 % coverage) separated the larger flavophages into two groups (Fig. S2): FKj−2, FKy−2, and FKo−2 (∼200 kb) formed one clade, while FKo−1 and FTs−1 (∼390 kb) grouped together on a distinct branch. VIRIDIC results mirrored this pattern, showing high relatedness only within these two clusters, with all other jumbophage pairs sharing <10 % intergenomic similarity (Supplementary material, Fig. S3). Larger phages also clustered separately from smaller isolates, with only sparse PC matches between them. These patterns indicate no evidence of recent genome expansion and instead support independent evolutionary lineages associated with different genome size groups.

**Figure 2.**
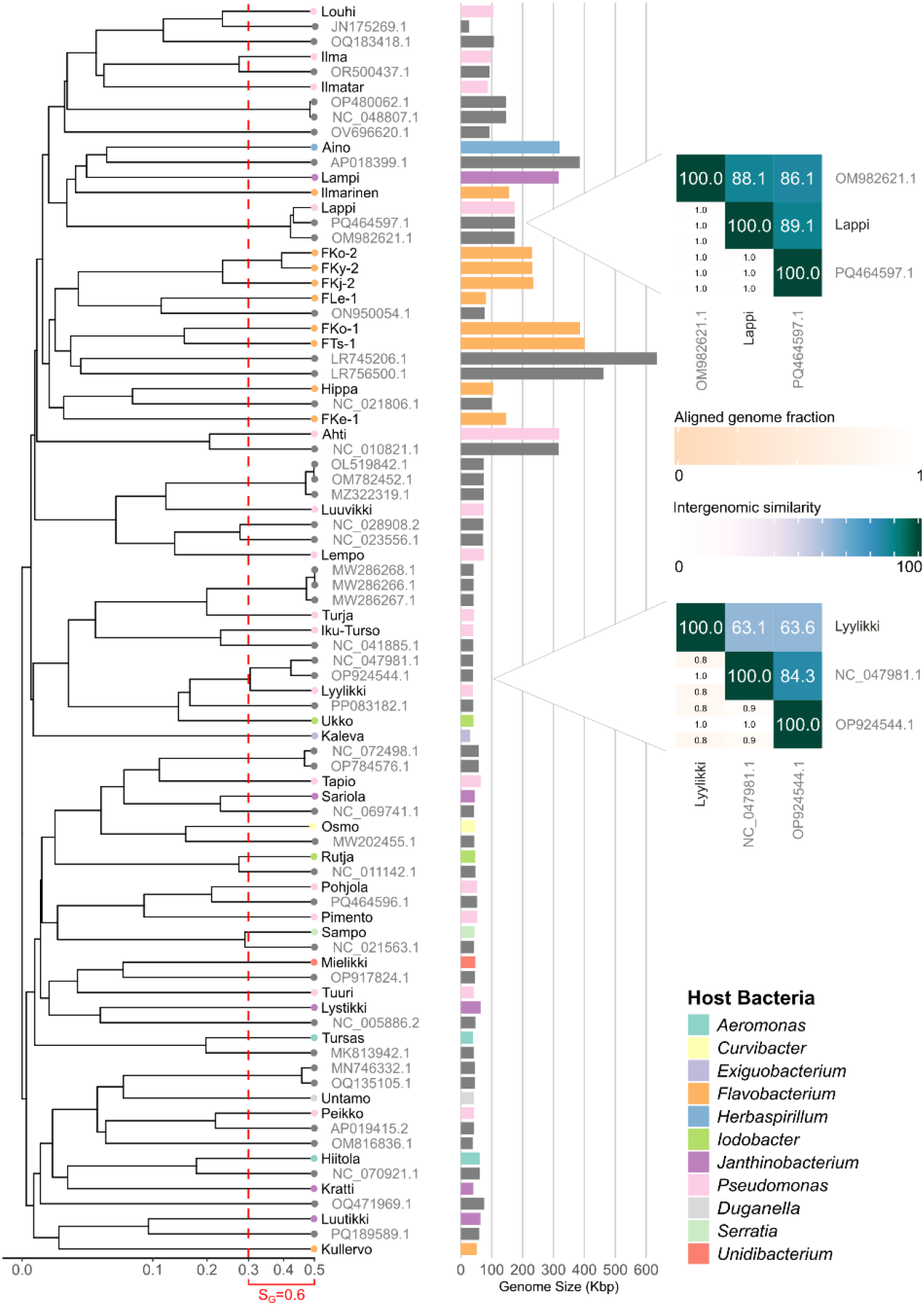
Comparison of the freshwater phages with selected reference genomes. On the left, a rectangular proteomic tree generated with the ViPTree online server is shown. Names of freshwater phages are in black, while accession numbers of reference phages retrieved from the INPHARED database are in grey. The tree was calculated from genome-wide distances based on normalized tBLASTx scores and was rooted at the midpoint. A red dashed line indicates a genomic similarity (S_G_) threshold of 0.6, used to delineate subgroups of closest relatives. On the right, a bar plot displays phage genome lengths, colour-coded by the genus of the bacterial host used for isolation (colour scheme shown in the legend, bottom right) with bars representing reference phages shown in grey. For the closest phage representatives, VIRIDIC results for Lappi and Lyytikki are provided as footnotes. Heatmaps show aligned genome fractions and intergenomic similarities for the VIRIDIC analysis.

### Freshwater phages represent novel genomic diversity

Whole-genome BLAST searches against the INPHARED database revealed no species-level relatives for any of the 40 phages. For each phage, the top-ranked genome match with the highest query coverage was selected for further analysis when available. A proteomic tree was subsequently constructed using ViPTree, including the 40 phages from this study alongside 43 reference genomes (Fig. 2). To refine the comparison for the most similar phages, they were compared using VIRIDIC. At the genus level, relatedness to previously described phages was revealed only for Lappi infecting *Pseudomonas*. It showed similarity to *Pseudomonas* phages Milchi (PQ464597.1) and Astolliot (OM982621.1), with intergenomic similarities of 89 % and 88 %, respectively. Both of the matching phages are currently unassigned to any genus. All other phages lacked close matches, underscoring the high diversity of obtained freshwater isolates.

Phage genomes ranged from 30 kb to 400 kb (Fig. 1). Among the 40 genomes, 27 were below 100 kb, five ranged between 100–200 kb, and eight exceeded 200 kb, thus qualifying as jumbophages (Yuan & Gao, 2017). The unexpectedly high proportion of jumbophages is notable given that all phages were obtained using conventional enrichment methods with endogenous freshwater bacteria. Five of the eight jumbophages infected *Flavobacterium* hosts, with the remaining three infecting *Pseudomonas, Janthinobacterium*, and *Herbaspirillum*. To our knowledge, these include the first reported jumbophages infecting *Janthinobacterium* and *Herbaspirillum*.

### Jumbophages display large myovirus morphology

Transmission electron microscopy revealed that all jumbophages exhibited myovirus morphology, with isometric capsids and long contractile tails (Fig. 3, S4, see also Fig. S5 for non-jumbophage isolates). FKo-2 has previously been imaged (Laanto et al., 2011). Capsid diameters (facet-to-facet) ranged from 115 to 149 nm (mean = 131 nm), while tail lengths varied from 112 to 187 nm (mean = 157 nm) (Table S6). Janthinobacterium infecting phage Lampi had the largest virion with a capsid diameter of 146 nm and a tail length of 141 nm. Overall, morphology was consistent across isolates, suggesting a shared structural architecture despite their genomic diversity (see below).

**Figure 3.**
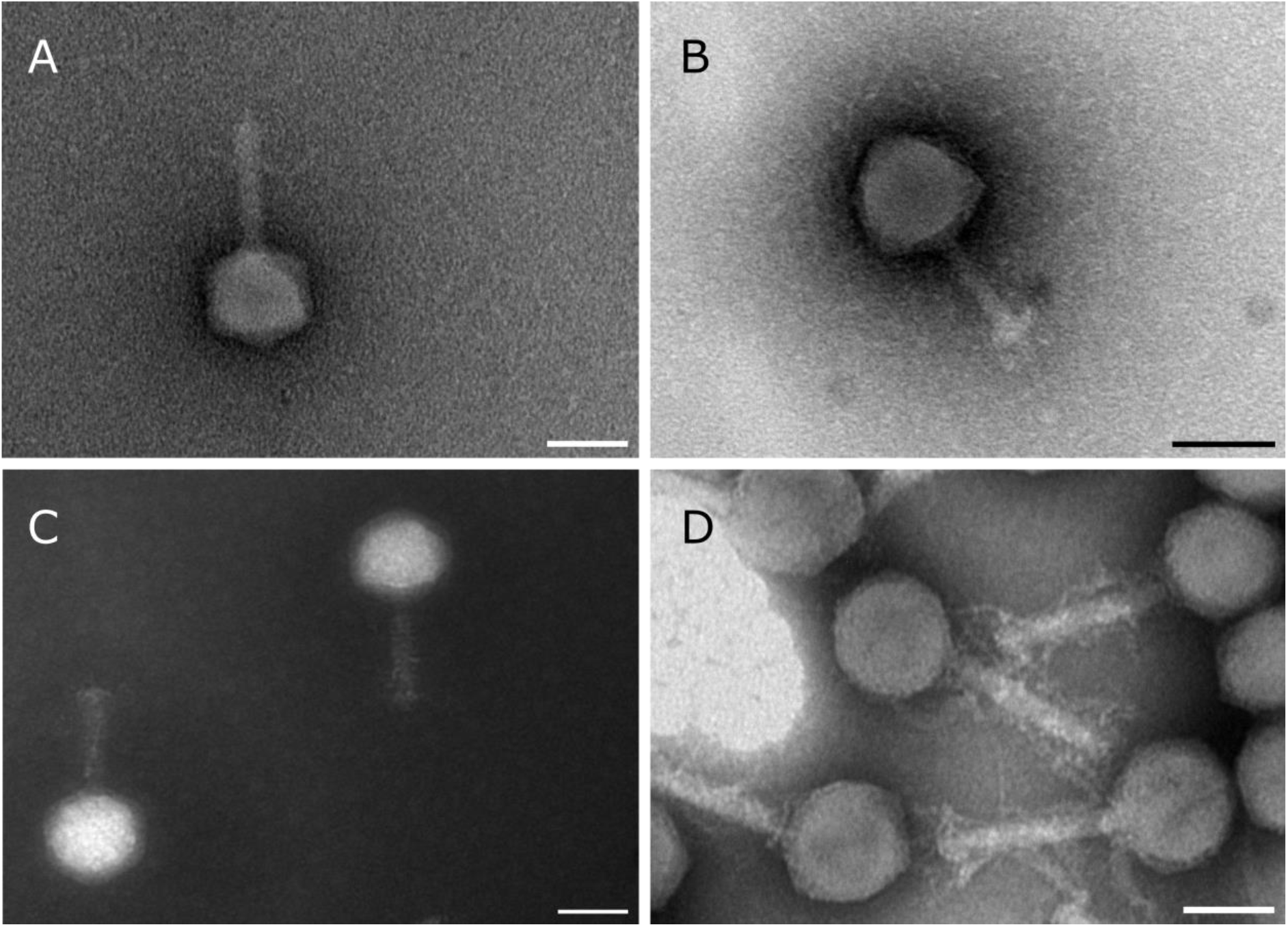
Transmission electron microscopy of negatively stained jumbophages. A) *Flavobacterium* phage FKo-1. B) *Herbaspirillum* phage Aino. C) *Janthinobacterium* phage Lampi. D) *Pseudomonas* phage Ahti. Scale bars: 100 nm.

### Jumbophage–host interactions

For most jumbophage–host pairs, low-MOI infections (0.01–0.1) resulted in growth curves indistinguishable from uninfected controls, suggesting limited infection efficiency at low initial phage-to-cell ratios (Fig. 4). In contrast, several phages exhibited clear inhibitory and lytic effects, such as a reduced growth rate, at MOI 1. Overall, these results indicate a strong MOI dependence in detectable culture-level effects. This pattern is consistent with the longer latent periods reported for some jumbophages (Sharma et al., 2019; Ahmad et al., 2021; Hu et al., 2023).

**Figure 4.**
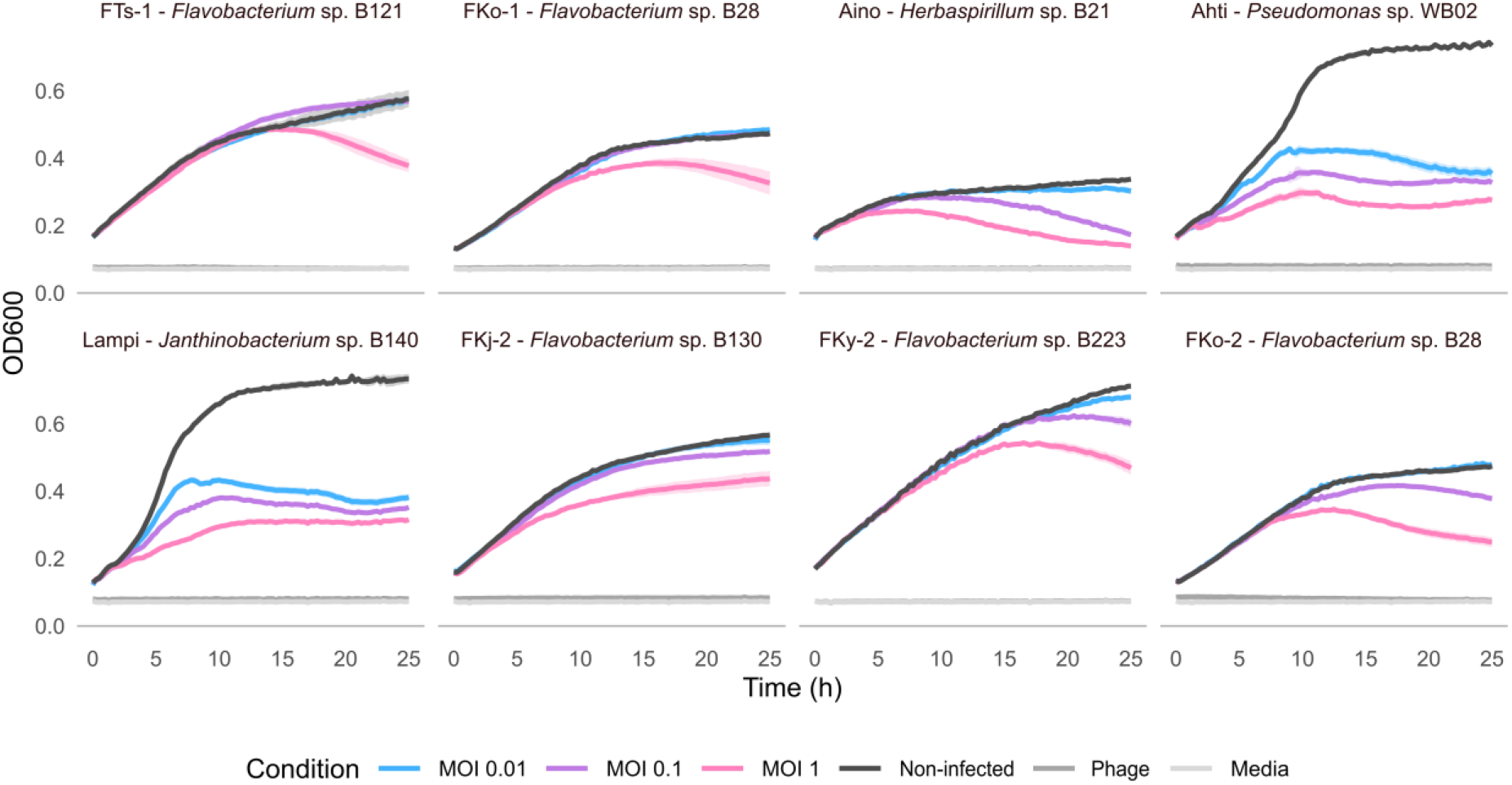
Jumbophage infections. Optical density (OD_600_) measurements of host cultures infected with each jumbophage at three MOIs (0.01, 0.1, and 1). Each host–phage panel represents an individual phage isolate, with its host strain. Curves represent the mean OD_600_ of five replicate wells, and shading indicates ±SD.

Host range assays conducted across all jumbophage hosts showed that all jumbophages infected only bacterial strains belonging to the same genus as their original isolation host, indicating a narrow host range (Fig. S6). In addition, when tested against 12 *Pseudomonas* sp. strains, Ahti infected two additional strains. On their isolation hosts, the jumbophages predominantly formed small plaques (Fig. 5). *Herbaspirillum* phage Aino formed particularly small, turbid plaques, consistent with low infection efficiency or incomplete lysis (Fig. 4), which made quantitative assessment of plaques difficult.

**Figure 5.**
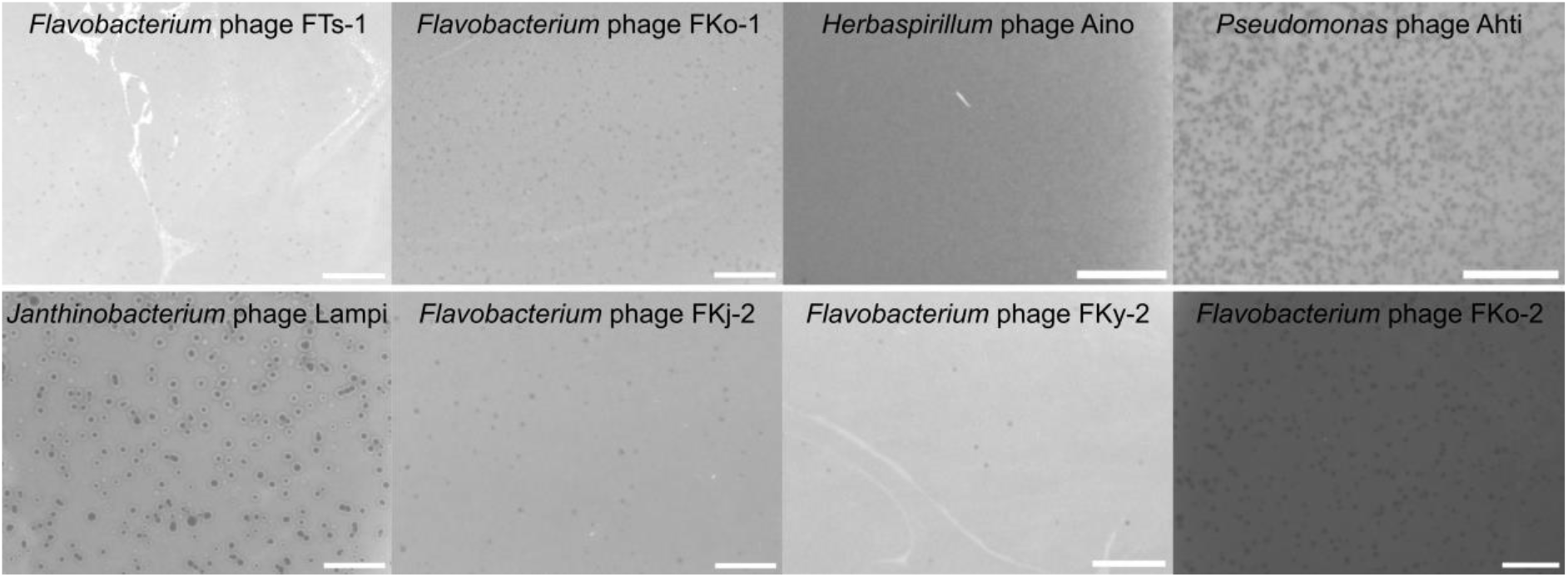
Plaque morphologies of the jumbophages on their isolation host bacteria at 24–48 h post-infection. Scale bars: 5 mm.

Phylogenetic analysis based on the large terminase subunit positioned the newly isolated jumbophages among established dsDNA phage lineages (Fig. 6, see also Fig. S7 for major capsid protein phylogeny). The eight new jumbophages did not form a single monophyletic cluster but instead grouped separately with representatives of multiple phage lineages. The *Flavobacterium*-infecting phages FKo-2, FKy-2, and FKj-2 clustered closely together, consistent with their higher intergenomic similarity values. In contrast, *Janthinobacterium* phage Lampi and *Herbaspirillum* phage Aino were positioned in more distant branches. Bootstrap support was high (>70) for most major nodes, confirming the reliability of the observed relationships. Collectively, the tree highlights the phylogenetic diversity of the jumbophages and their distribution across multiple branches, consistent with the broader polyphyletic nature of jumbophages reported previously (Yuan & Gao, 2017; Iyer et al., 2021).

**Figure 6.**
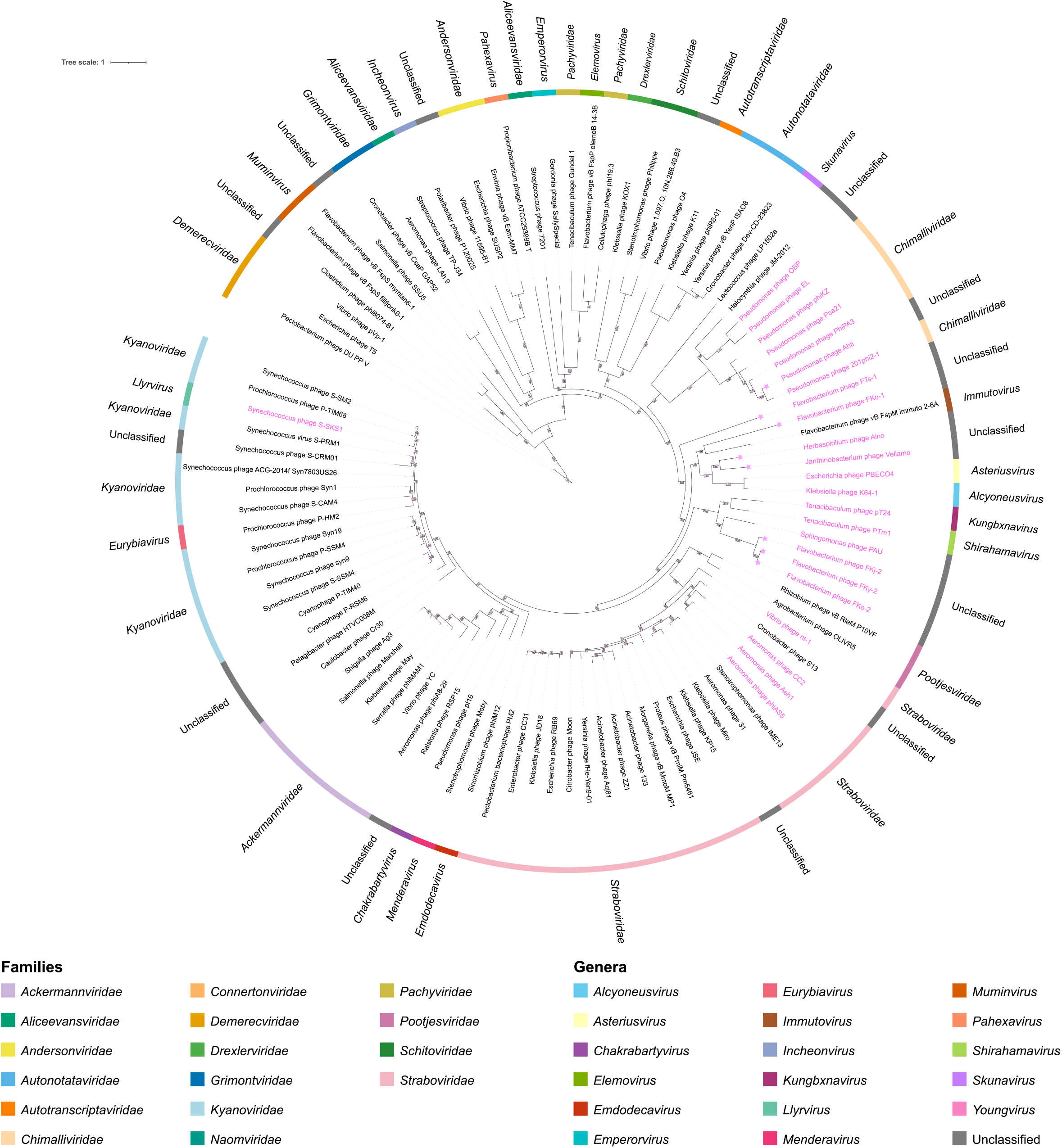
Maximum likelihood phylogenetic tree of large terminase subunit protein sequences from the eight freshwater jumbophages (stars) and 97 representative reference sequences from diverse tailed dsDNA phage lineages. Jumbophages are marked in pink. Analyses were performed using IQ-TREE with the best-fit evolutionary model selected by model testing and 1,000 bootstrap replicates. Bootstrap values are indicated for each node. The tree is midpoint-rooted and visualised in iTOL v7. The outer ring is colour-coded by phage family or genus (when not assigned to a family).

### Functional annotation highlights diverse biological capacities

Genome annotation revealed substantial variation in functional content across the phage collection, with larger genomes containing a higher proportion of unannotated genes and more putative metabolic- and host-interaction–related functions than the smaller ones (Supplementary material, Fig. S8). The *Pseudomonas* jumbophage Ahti uniquely encoded numerous head-and-packaging genes, while the largest genomes showed increased COG-level matches, including a ribosomal protein in Aino. Given that larger phage genomes are known to encode more tRNAs (Bailly-Bechet et al., 2007; Al-Shayeb et al., 2020), we examined tRNA gene content within our collection. Predicted tRNA counts ranged from 0 to 62 and tended to increase with genome size, yet several of the largest jumbophages encoded only a few tRNAs or none at all (Supplementary material, Fig. S9).

Across the collection, we identified genes representing all major functional categories implicated in the control and modification of host metabolism (Table S3 & S4). A broad array of genes associated with transcription and translation was present in all genomes. The phages Luuvikki and Lempo each encoded a putative virion RNA polymerase, while the two largest jumbophages, FKo-1 and FTs-1, carried two distinct RNA polymerase σ-factor variants per genome. In the *Flavobacterium* phage Hippa, a large, predicted protein (2,175 residues; gene V2_00052) contained the conserved DFDID motif, characteristic of distant RNA polymerases previously described in crAss-like phages (Drobysheva et al., 2021).

Several large genomes also encoded tRNA synthetases and proteins involved in tRNA modification or ligation, whereas a translation initiation factor was predicted exclusively in the largest FTs-1 genome. Genes related to host cell lysis were detected in all phages except Tursas and Kullervo. In contrast, genes potentially associated with temperate replication were restricted to smaller phages. Notably, an “excisionase and transcriptional regulator” was predicted only for the smallest phage in the collection, Kaleva.

### Metabolic potential of the jumbophage isolates: candidates for AMGs

We identified putative AMGs from jumbophages (Table S7). A phage-encoded enzyme, a nicotinamide phosphoribosyltransferase (NAMPT), not previously detected in jumbophages, was present in the largest phages Ahti, Aino, FTs-1, and FKo-1. This enzyme, part of the NAD^+^ salvage pathway (Zhang et al., 2025), has previously been reported in Vibrio phage KVP40 (Lee et al., 2017). In the *Flavobacterium* jumbophages FTs-1 and FKo-1, the NAMPT gene occurs within a ∼10 kb long conserved gene cluster in which other genes related to NAD–salvage were also detected (Fig. 7, Table S8). Here, we refer to these clusters as phage NAD–salvage islands. For both *Flavobacterium* jumbophages, the islands include a gene encoding a protein with a likely bifunctional architecture: an N-terminal nucleotidyltransferase (NMNAT-like) domain and a C-terminal Nudix hydrolase domain. This gene is located near the NAMPT gene, indicating operon-level co-regulation.

**Figure 7.**
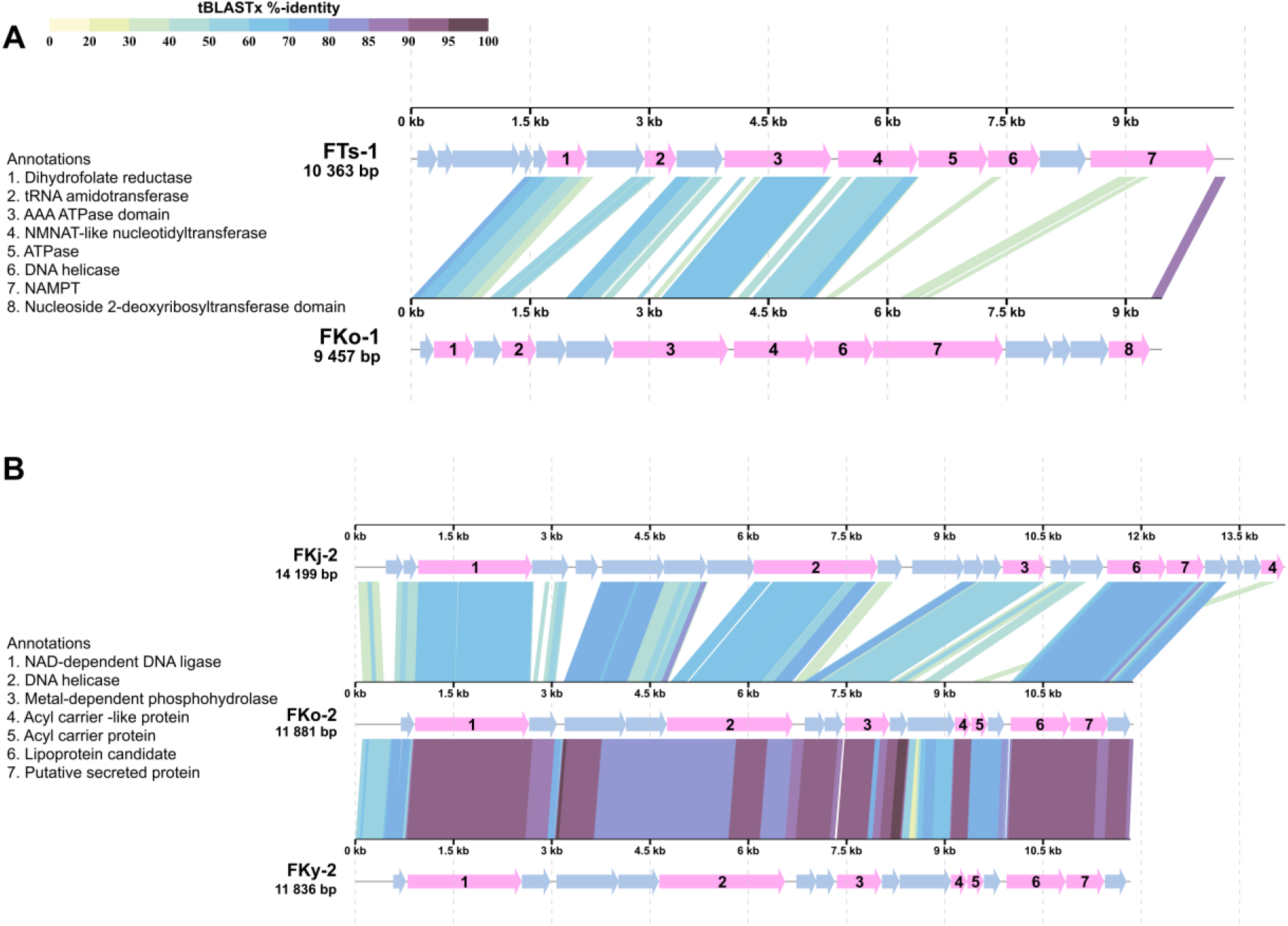
AMG islands in freshwater jumbophages. Alignments highlighting the two major putative AMG modules identified in the jumbophage genomes. A) NAD–salvage module in jumbophages FTs-1 and FKo-1. B) Host-interface island in jumbophages FKj-2, FKo-2, and FKy-2. Coloured ribbons indicate homologous proteins (tBLASTx percent identity). Genes with functional annotations are highlighted in pink.

Another distinct AMG module was identified in the *Flavobacterium* phages FKo-2, FKj-2, and FKy-2 (Fig. 7, Table S8) that we named Host-interface island. For example, in FKo-2 we identified a 17-gene island with secreted and lipoprotein candidates, duplicated acyl-carrier proteins, and a metal-dependent phosphohydrolase. This gene composition suggests a phage-encoded host-surface remodelling module likely involved in adsorption or envelope modification. In FKj-2, the annotated acyl carrier proteins differ at the nucleotide level from those of the other two phages, but the overall proximity of genes is retained. While phage-encoded ACPs and secreted effectors have been reported in other jumbophages (Kim et al., 2020), the recurrent and syntenic occurrence of this multi-gene module across different genomes suggests it represents a conserved accessory strategy for host-surface modulation in freshwater flavobacterial phages. Thus, we suggest this to be a putative phage-encoded module that coordinates targeted acylation of membrane proteins.

### Anti-defence systems are prevalent in jumbophages

Genomic analyses revealed multiple putative anti-defence systems, with detection varying slightly depending on the annotation method used (Phagenomics versus Pharokka + Phold) (Table S9). Among the non-jumbophages, eight out of 32 genomes contained at least one predicted anti-defence element. In contrast, a higher number of jumbophages had predicted anti-defence elements. Five of the eight jumbophages contained at least one system, with several having two identified systems. Within the jumbophages, Anti-Thoeris elements were the most frequently identified, predicted in four genomes (FKo-1, FTs-1, Lampi, and Ahti) by at least one annotation-tool combination. Anti-CBASS systems were found in FKo-1 and FKy-2, and Anti-Gabija was detected only in Aino. Additionally, one putative system, NADP, was found in Ahti. PadLoc analyses further revealed PDC-S10 systems in FKo-1 and FTs-1, confirmed by both annotation pipelines.

No CRISPR arrays or *cas* genes were found in any of the jumbophage genomes. Comparative searches for the core jumbophage genes defined by Iyer et al. (2021) showed that most were absent from the jumbophage genomes described here, with only a small subset of replication- and repair-associated homologues detected (Fig. S10). These findings suggest that the freshwater jumbophages possess distinct functional repertoires compared with previously characterised marine and *Pseudomonas* jumbophages (Iyer et al., 2021).

### Pseudomonas jumbophage Ahti is a freshwater nucleus-forming phage

During infection of *Pseudomonas* sp. WB02 by phage Ahti, confocal microscopy revealed distinct localisation of DAPI-stained DNA within host cells, in contrast to the diffuse staining pattern observed in the uninfected control (Fig. 8A–B). This indicates compartmentalisation of phage DNA during infection, consistent with the formation of a nucleus-like structure (Laughlin et al., 2022). No comparable localisation was detected for any other freshwater jumbophage infection (Fig. S11). Genomic analysis revealed that Ahti encodes homologues of predicted tubulin and chimallin proteins which are associated with known nucleus-forming phages (Chaikeeratisak et al., 2021; Prichard et al., 2023). Furthermore, Ahti contained homologues for all 21 core genes defined as exclusive to nucleus-forming phages of the family *Chimalliviridae* (Prichard et al., 2023), most of which are annotated as hypothetical proteins (Fig. 8C & S12). The strongest matches were observed between Ahti and ΦKZ, with weaker similarity to Goslar and RAY, consistent with *Pseudomonas* phage ΦKZ being more similar to Ahti. Terminase-based phylogeny further placed Ahti within *Chimalliviridae* (Fig. 6). Together, these findings indicate that Ahti represents a new member of the family *Chimalliviridae* and is, to our knowledge, the first nucleus-forming freshwater jumbophage.

**Figure 8.**
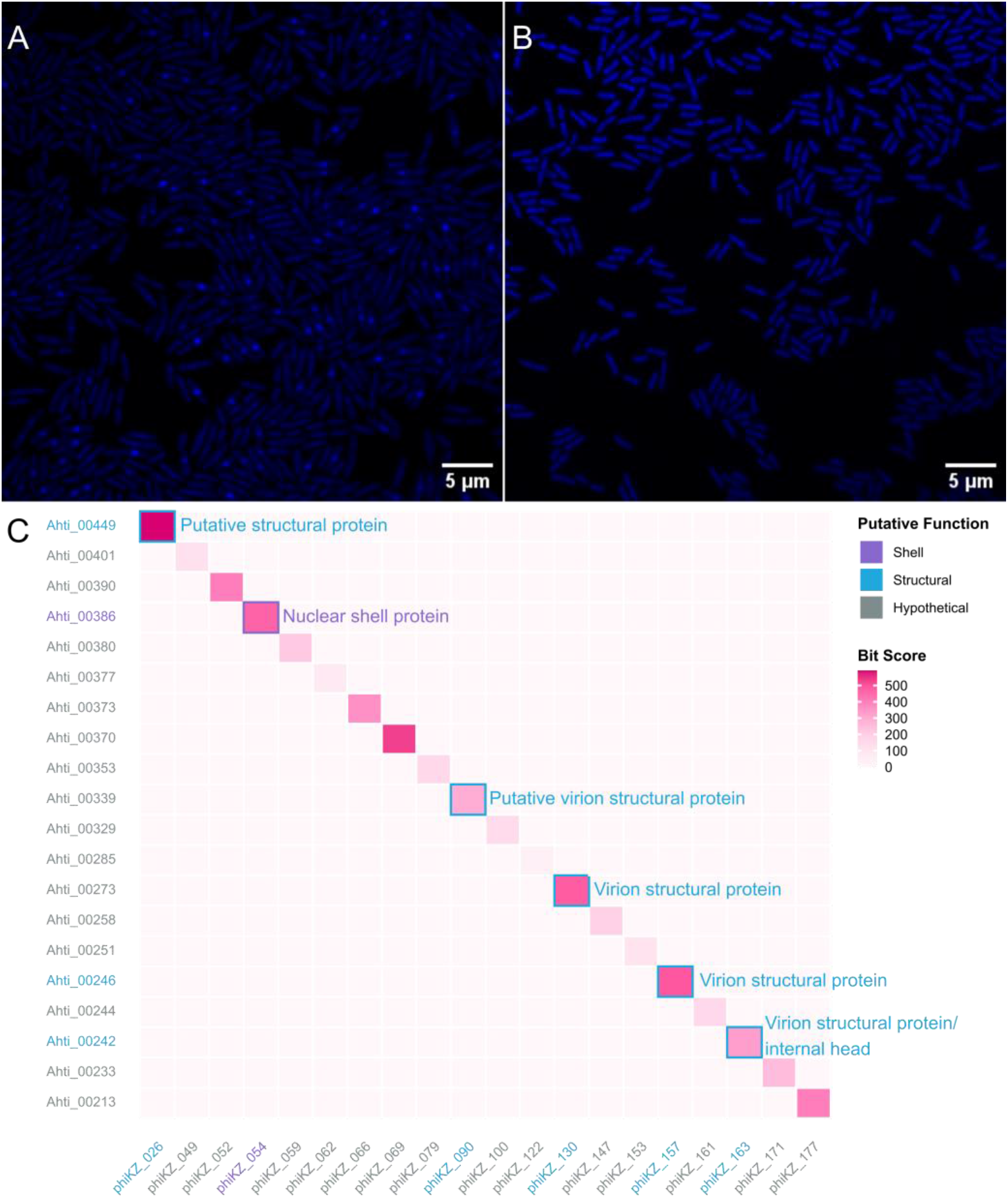
Nucleus-like compartment formation and presence of nucleus-forming core gene products in phage Ahti. A) Confocal microscopy of *Pseudomonas* sp. WB02 infected with phage Ahti at 60 min post-infection showing distinct localisation of DAPI-stained DNA. B) Uninfected control cells displaying diffuse DAPI staining. C) Heatmap of BLASTp bit scores (0–500) comparing the Ahti proteome to the 21 core gene products defined as exclusive to nucleus-forming phages (Prichard et al., 2023). ΦKZ was used as the reference genome for these core genes due to its genetic similarity to Ahti. Because ΦKZ lacks one core gene and contains three paralogous copies of another (for which only the highest-scoring match is shown), the heatmap displays 20 unique core-gene matches. Gene products are colour-coded by predicted functional class, and text labels indicate putative functions of the corresponding ΦKZ genes (Prichard et al., 2023).

## Discussion

Metagenomic surveys consistently highlight the immense viral diversity of aquatic systems (López-Pérez et al., 2017; Coutinho et al., 2018; Kavagutti et al., 2019). However, isolate-based studies remain essential to validate phage–host interactions and test predicted functions experimentally. While 80 % of metagenomics datasets are derived from environmental samples (IMG/VR database) and marine environments are recognised as hotspots of phage diversity (Suttle, 2005), phage banks are dominated by soil- and clinic-derived isolates. Systematic collections of environmental isolates remain scarce. By isolating diverse endogenous phage– host pairs from freshwaters, we established a unique collection that reflects ecologically relevant interactions. Our results show that boreal and subarctic freshwater environments serve as an underexplored reservoir of unique non-model systems.

In total, 40 unique phages were isolated infecting bacterial hosts from 11 phylogenetically diverse genera. Surprisingly, an unexpectedly high proportion of jumbophages was detected, despite relying on conventional enrichment-based isolation methods and endogenous host bacteria. These jumbophages span diverse evolutionary lineages, including the first known jumbophage isolates infecting *Herbaspirillum* and *Janthinobacterium*. In addition, we describe one *Pseudomonas* jumbophage and five *Flavobacterium* jumbophages. Jumbophages showed no evidence of a single origin. Instead, they formed distinct lineages, including two divergent *Flavobacterium-*infecting groups with genome sizes of approximately 235 kb and 390 kb. These patterns reflect long-term evolutionary separation and high genomic diversity.

Although unusual among host-dependent viruses, large genomes are also found in giant eukaryotic dsDNA viruses of the phylum *Nucleocytoviricota*, which are important ecological players in cold freshwater environments and can reach several megabases in size (Ha et al., 2021; Zhang et al., 2024). In jumbophages, expanded genome size may support broader functional capacity and reduce reliance on host metabolic processes (Yuan & Gao, 2017). Jumbophages can fine-tune host resource utilisation through AMGs. We identified two putative AMG islands in *Flavobacterium*-infecting jumbophages, one centred on NAD–salvage and another on putatively modifying the host cell surface or envelope. The NAD–salvage island can aid in viral replication by boosting NAD^+^ availability. In turn, the Host-interface island putatively supports host biosynthetic activity during infection. Their presence across jumbophages suggests selective advantages in nutrient-limited and variable freshwater environments. These islands may represent transferable functional modules shaping phage–host interactions in lakes and rivers. While marine and *Pseudomonas* jumbophages have been shown to encode certain core functions and adaptive features (Iyer et al., 2021), the freshwater phages stand out, since they lack homologues to most of these genes. These findings emphasise that jumbophages do not share a standard functional repertoire but instead employ diverse genomic strategies to thrive in different ecological contexts.

A larger genome may also confer jumbophages with more efficient strategies for host takeover. In our jumbophages, we detected transcription- and translation-related proteins, and a diverse repertoire of predicted tRNAs, which may also increase replication efficiency (Nazir et al., 2021). Moreover, recent bioinformatic evidence indicates that some phages even encode ribosomal proteins (Mizuno et al., 2019), potentially representing a novel strategy for manipulating host translation. Their functionality, however, remains experimentally unvalidated (Mizuno et al., 2019). Notably, within our collection, *Herbaspirillum* phage Aino was predicted to encode a ribosomal protein L7/L12 homologue, highlighting Aino’s potential as an experimental model for assessing whether such genomic predictions reflect real biological function.

Host defence systems can be detrimental to phage propagation (Egido et al., 2021). Thus, another way for jumbophages to benefit from their increased genomic capacity is to ensure a successful infection despite these defences. A variety of anti-defence systems have been described for phages (Yirmiya et al., 2024), and in our collection, a proportionally higher number of jumbophages had predicted anti-defence elements compared to the smaller phages. Given the high proportion of unannotated genes among large genomes, the likelihood to discover yet-uncharacterised systems remains high. Jumbophage genomes can also encode more complex infection strategies to evade host immunity, such as forming a protective phage nucleus. This feature distinguishes members of the family *Chimalliviridae* (Prichard et al., 2023). In our study, *Pseudomonas* phage Ahti exhibited this strategy, as confirmed by confocal microscopy. Genomic analysis further supported this observation: Ahti harbours a tubulin gene as well as homologues to all 21 core genes defined for *Chimalliviridae* (Prichard et al., 2023). Furthermore, Ahti clusters with known members of this family in terminase-based phylogenetic analysis. We therefore propose Ahti as a new species of the family *Chimalliviridae*, representing, to our knowledge, the first nucleus-forming jumbophage isolated from freshwater.

Overall, our study underscores boreal and subarctic freshwaters as rich reservoirs of non-model bacterium–phage systems, revealing novel jumbophage lineages, new host associations, and diverse biological strategies. Described isolates provide cultured representatives of the viral ‘dark matter’ frequently observed in metagenomic surveys transforming sequence-based predictions into tangible biological entities. This jumbophage collection offers valuable experimental model systems to challenge the traditional boundary between cellular and non-cellular life. By expanding the taxonomic spectrum of known phages, these cultured isolates offer a foundation for investigating how jumbophages shape freshwater microbiomes and drive ecosystem dynamics.

## Acknowledgements

The authors want to thank Sari Korhonen, Tero Sievänen, Johannes Nummela, Irini Assimakopoulou, Diego Gonzalez Martin-Calero, Juliaana Kauraala, Janita Nieminen, and Janika Bergfors for technical assistance. In addition, we thank Visa Ruokolainen and Petri Papponen for valuable help with confocal and electron microscopy. We also acknowledge everyone who contributed to the sample collection. The facilities and expertise of the Instruct-HiLIFE Biocomplex unit at the University of Helsinki, a member of Instruct-ERIC Centre Finland, FINStruct, and Biocenter Finland are gratefully acknowledged. We made use of geospatial data provided by the Open Geospatial Information Infrastructure for Research (Geoportti, urn:nbn:fi:research-infras-2016072513) funded by the Research Council of Finland, CSC – IT Center for Science, and other Geoportti consortium members. Large language models (ChatGPT 5.1, OpenAI) were used to assist with code generation and text editing during manuscript preparation. All AI-assisted content was reviewed, revised, and approved by the authors, who take full responsibility for the final manuscript.

## Author contributions

Conceived the study: EL, LRS, HMO; sampling: EL; experimental work: MN, HMO, EL; data analysis: MN, KK, HMO, EL; writing – first draft: MN, KK; writing – review and commenting: MN, KK, LRS, HMO, EL; supervision: LRS, HMO, EL; resources: LRS, HMO, EL

## Data availability

For review purposes, data can be found from the link provided. Upon final submission, all data will be made openly available in accordance with journal policy.

